# Heat shock-mediated transformation is possible in several Gram-negative bacteria

**DOI:** 10.1101/2025.10.30.685485

**Authors:** Shubhra Jyoti Giri, Pankaj Losan Sharma, Lukapriya Dutta, Shuhada Begum, Shuvam Bhuyan, Monika Jain, Niraj Agarwala, Suvendra Kumar Ray

## Abstract

Transformation of foreign DNA, subjecting *Escherichia coli* to CaCl_2_ treatment followed by heat shock exposure, is a regular approach in recombinant DNA technology. However, in spite of its large popularity in *E. coli*, heat shock transformation is rarely reported in other Gram-negative bacteria. Instead, techniques such as natural transformation, conjugation, or electroporation are used in those bacteria. In this study, we have successfully implemented the heat shock transformation in four different Gram-negative bacteria, such as *Ralstonia pseudosolanacearum, Pseudomonas aeruginosa, Pseudomonas putida*, and *Enterobacter roggenkampii*. The standard heat shock transformation procedure used for *E. coli* DH5α has been modified. The pDSK-GFPuv plasmid bearing the green fluorescence gene was directly transferred to *Ralstonia pseudosolanacearum* by heat shock at 50 ºC for 60 seconds. For *Pseudomonas aeruginosa, Pseudomonas putida*, and *Enterobacter roggenkampii*, the cells were made competent using CaCl_2,_ followed by performing transformation by heat shock at 50 ºC for 180 seconds. The transformants were resistant to kanamycin as well as exhibited fluorescence. These transformants were used to study the colonization pattern of tomato seedlings. Our study suggested that the heat shock transformation method can also be performed to introduce genes in other Gram-negative bacteria, other than *E. coli*.

## 1 Introduction

Bacterial transformation is an important process of horizontal gene transfer [1,2] in both Gram-positive and Gram-negative bacteria. Several bacteria are known to exhibit their natural ability to uptake foreign DNA from the external medium through natural transformation. This ability has been exploited by researchers to introduce foreign DNA into these bacteria for different research purposes [3-5].

Due to the limitation of natural competence in *Escherichia coli*, artificial transformation is popular in this bacterium, that involves making bacterial cells competent or susceptible under controlled laboratory conditions to facilitate DNA uptake. In this process, the competence of bacteria is induced mainly by treating the cells with CaCl_2,_ followed by heat shock treatment. The Ca^2+^-rich environment not only helps overcome the repulsion between the external DNA and bacterial membrane owing to negative charges on them both but also creates a static force of attraction within the external DNA molecule that facilitates its folding into a compact ball-like structure [6-8]. Following the CaCl_2_ exposure, a quick temperature rise is applied to the bacterial cells, which results in a heat shock. This heat shock causes pores or gaps to form temporarily in the bacterial membrane. These openings facilitate increased permeability of the membrane, making it easier for the condensed DNA to be transferred into the bacterial cell. The heat shock also affects plasmid supercoiling, which might add to the transformation ability [9, 10]. Because of its easy nature, heat shock is a widely used method to do transformation in *E. coli*. However, due to certain limitations, such as the introduction of large DNA molecules by this method, the electroporation technique has been widely used for many Gram-negative and Gram-positive bacteria. In electroporation transformation, a special electroporator device is required through which an electric pulse is applied to create a pore in the bacterial membrane. Because of the high electric voltage used during electroporation, there are certain disadvantages to using this technique in bacteria [11].

The use of heat shock transformation has not been reported widely for other bacteria except *Escherichia coli*. Our laboratory is working with several Gram-negative bacteria, such as *Ralstonia pseudosolanacearum*, the causal agent of the bacterial wilt disease in plants, and plant-associated bacteria such as *Pseudomonas aeruginosa, Pseudomonas putida*, and *Enterobacter roggenkampii*. These three plant-associated bacteria isolated from tomato seedlings inhibit the phytopathogen *R. pseudosolanacearum* (unpublished results). We want to transfer the GFP plasmid to these four Gram-negative bacteria by heat shock-mediated transformation. A green fluorescent plasmid, pDSK-GFPuv having the size of ∼ 8.9 kb, was successfully transferred into these Gram-negative bacteria by heat shock-mediated transformation at different temperatures and times, respectively. After successfully transforming the GFP plasmid into the targeted bacterial strains, these were used in the study of the association of bacteria with plants. The introduced plasmid was re-isolated from one of the bacteria, *R. pseudosolanacearum* (TRS4000), to ensure the transformation of the GFP plasmid by the heat shock method. This is the first elaborate heat shock-mediated transformation (HSMT) in different Gram-negative bacteria. Our work indicates that different bacteria might be targeted through heat shock transformation.

## 2 Materials and methods

### 2.1 Chemicals and growth media

All chemicals, bacterial culture medium components, and antibiotics used in this study were procured from Hi-Media, Mumbai, India, and SRL, New Mumbai, India. Plastic wares were bought from Tarsons, Kolkata, India; Glassware was procured from Borosil, Kolkata, India.

### 2.2 Bacterial strains and growth conditions

*Ralstonia pseudosolanacearum* F1C1, *Pseudomonas aeruginosa* SPT08, *Pseudomonas putida* N4T, and *Enterobacter roggenkampii* SPT20 are isolated from natural sources in the laboratory. For this study, these bacteria were grown at 28 °C in BG medium, which contains 1% peptone, 0.1% yeast extract, and 0.1% casamino acids supplemented with 5 g/L glucose [12]. Solid agar medium was prepared using 1.5% agar. *E. coli* DH5α cells were grown in Luria–Bertani medium at 37 ºC [13]. Kanamycin antibiotic has been used at a concentration of 50 μg/mL. Bacterial strains and the plasmid used in this study are listed in Table 1.

**Table 1:**
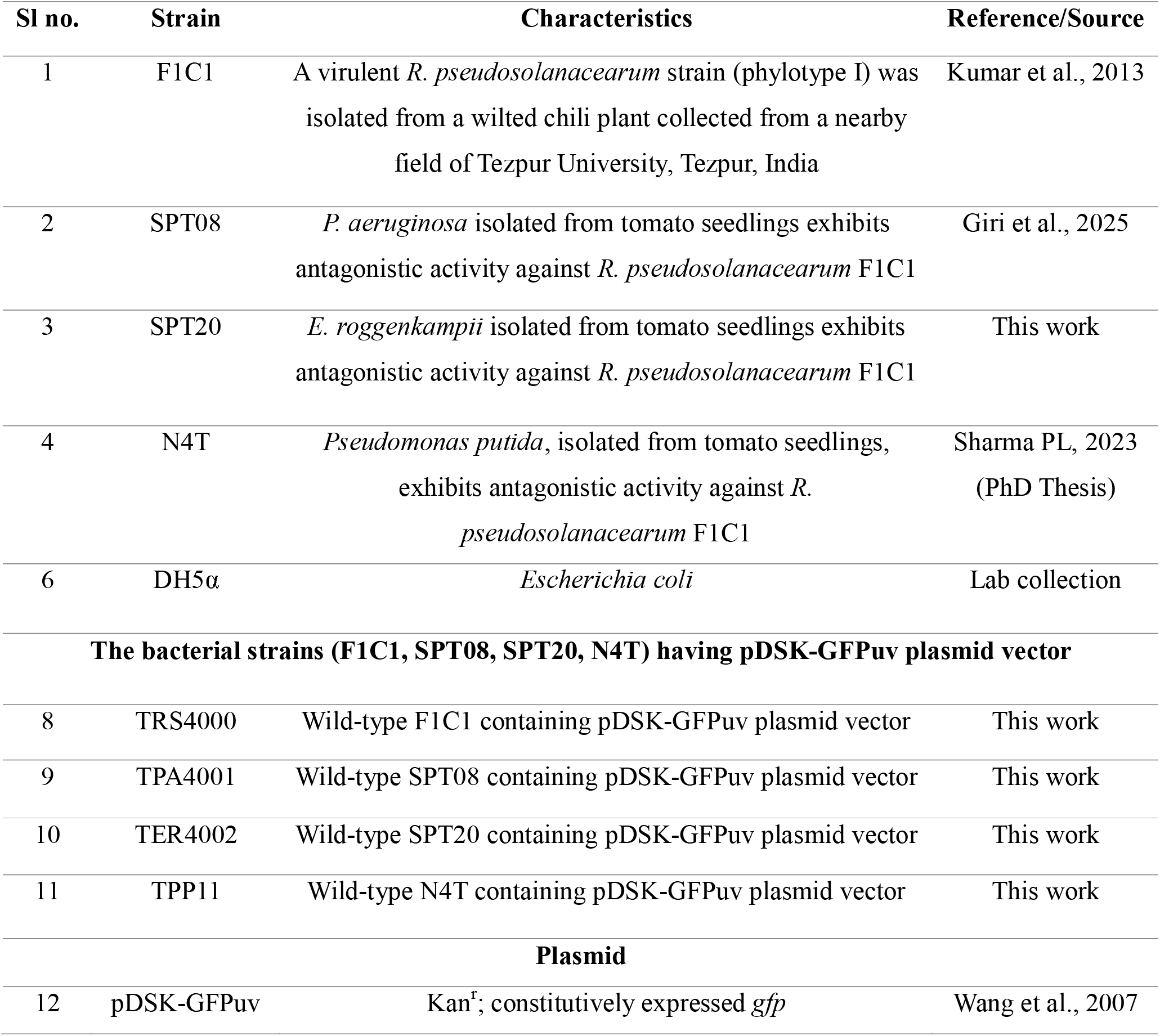
List of bacterial strains, plasmid vector used in this study.

### 2.3 Preparation of competent cells for *Pseudomonas aeruginosa, Pseudomonas putida*, and *Enterobacter roggenkampii*

Competent cells of *Pseudomonas aeruginosa, Pseudomonas putida*, and *Enterobacter roggenkampii* were prepared by inoculating the bacterial colony into BG liquid medium and allowing it to grow at 28 °C. A volume of 100 µL of saturated culture was inoculated in 10 mL of BG broth at 28 °C in a shaking incubator until it reached an absorbance of 0.5 at 600 nm. The cells were harvested by centrifugation at 4000 rpm for 10 minutes at 4 ºC. The cells were washed twice with 10 mL of 10 mM NaCl, harvested by centrifugation, and resuspended in 10 mL of 100 mM ice-cold CaCl_2._ The cells were kept on ice for 30 minutes, harvested by centrifugation (4000 rpm, 10 minutes, 4 °C), and re-suspended again with 100 mM ice-cold CaCl_2_ containing 15% glycerol. Five hundred microliters (500 µL) of the resulting harvested cells were distributed in 1.5 mL microfuge tubes and incubated for 12-24 h at 4 ºC, which increases cell competency. The aliquots were used for transformation and preserved in a deep freeze for future use [14, 15].

### 2.4 Heat shock transformation of pDSKGFPuv plasmid vector

The plasmid pDSK-GFPuv, a stable and broad host range plasmid containing one copy of constitutively expressed GFP and kanamycin resistance genes [16], was obtained from Dr. Kiran K. Mysore from the Plant Biology division, The Samuel Roberts Noble Foundation, USA. This plasmid was introduced to the prepared competent cells of *Pseudomonas* and *Enterobacter sp*. cells by the heat shock-mediated transformation method as described by Zhao et al. (2013) with a little bit of modification. Briefly, 50 µL of competent cells were mixed with 3 µL (1.5 µg/μL) of the pDSK-GFPuv plasmid. The mixture was then placed on ice for 30 minutes, followed by heat shock for 3 minutes at 50 ºC. After that, the tubes were immediately chilled in ice for 2 minutes and 30 seconds. After the treatment, 900 µL of BG broth was added to the mixture, and the cells were allowed to incubate for 6 h at 28 °C in a shaking incubator. The cells were pelleted down at 4000 rpm for 10 minutes at 4 ºC and resuspended in BG broth. A volume of 100 µL cell suspension was spread on a BG agar plate containing kanamycin antibiotic (50 µg/mL) as a selection marker. The successful transformants were further observed for the emission of green fluorescence under a fluorescence microscope (EVOS, Life Technologies) with 100X magnification.

For *R. pseudosolanacearum* F1C1, the heat shock-mediated transformation differs from that used for *Pseudomonas* sp. and *Enterobacter* sp. Cells of *R. pseudosolanacearum* were used directly for transformation, where the cells were grown in BG medium until their optical density (OD) reached 0.5 at 600 nm. A 50 µL volume of F1C1 cells was mixed with 3 µL (1.5 µg/µL) of the pDSK-GFPuv plasmid and incubated on ice for 30 minutes. The mixture then underwent a 60-second heat shock at 50 ºC, followed by immediate cooling on ice for 150 seconds. Afterward, 900 µL of BG broth was added, and the cells were incubated for 12 hours at 28 °C with shaking. The cells were harvested by centrifugation at 4000 rpm for 10 minutes at 4 ºC, resuspended in BG broth, and 100 µL of the suspension was spread onto a BG agar plate containing kanamycin. Plates were incubated at 28 ºC until colonies appeared. Single colonies were cultured in BG medium, and the micro-colonies of the transformants were examined for green fluorescence emission under a fluorescence microscope (EVOS, Life Technologies) at 100X magnification.

### 2.5 Confirmation of the GFP transformants

*R. pseudosolanacearum* was confirmed by phylotype-specific multiplex PCR [10, 18]. From the transformants, the green fluorescence plasmid (pDSK-GFPuv) was isolated and confirmed by restriction digestion with *EcoRI* to obtain the linearized plasmid. Furthermore, the GFP-tagged bacteria were confirmed by characterizing twitching motility, cellulase activity, and virulence assays, which were compared with those of the wild-type strains [19-24]. The transformants, *P. aeruginosa* SPT08 (TRS4001), *P. putida* N4T (TPP11), and *E. roggenkampii* SPT20 (TER4002), were checked for their antagonistic activity against *R. pseudosolanacearum*, as the wild type of these bacteria inhibits *R. pseudosolanacearum* F1C1.

### 2.6 Application of the GFP-tagged bacteria to study the plant-microbe association

Green fluorescence-tagged bacteria were further used in the study of plant-microbe association. To study the association with plants, seven-day-old tomato seedlings were used, which were grown in a sterile and controlled environment under a growth chamber (Orbitek, Scigenic). The seedlings were inoculated with each of the GFP-tagged bacteria via root inoculation. The seedlings were maintained in a growth chamber at 28 ºC, 75% relative humidity, and a photoperiod of 12 hours [22-25]. From the next day onwards of inoculation, the colonization of bacteria inside the seedlings was studied. Every day few seedlings were randomly selected and subjected to microscopic visualization. The seedlings were surface-disinfected as described by Kabyashree et al. (2020) and observed under a fluorescence microscope (EVOS, Life Technologies) with 100X magnification.

## 3 Results

### 3.1 Heat shock-mediated transformation of pDSK-GFPuv plasmid in Gram-negative bacteria

The plasmid, pDSK-GFPuv, carrying green fluorescent protein, was transformed into four plant-associated bacteria, *R. pseudosolanacearum* F1C1 (TRS4000), *P. aeruginosa* SPT08 (TPA4001), *P. putida* N4T (TPP11), and *Enterobacter roggenkampii* SPT20 (TER4002), by using the heat shock transformation method. The transformants were selected through a BG agar plate, containing kanamycin as a selection marker. The transformant colonies were observed under a fluorescence microscope using a GFP filter, and the expression of green fluorescence signal confirmed the successful transformation of the plasmid (Fig. 1A to D).

**Fig. 1.**
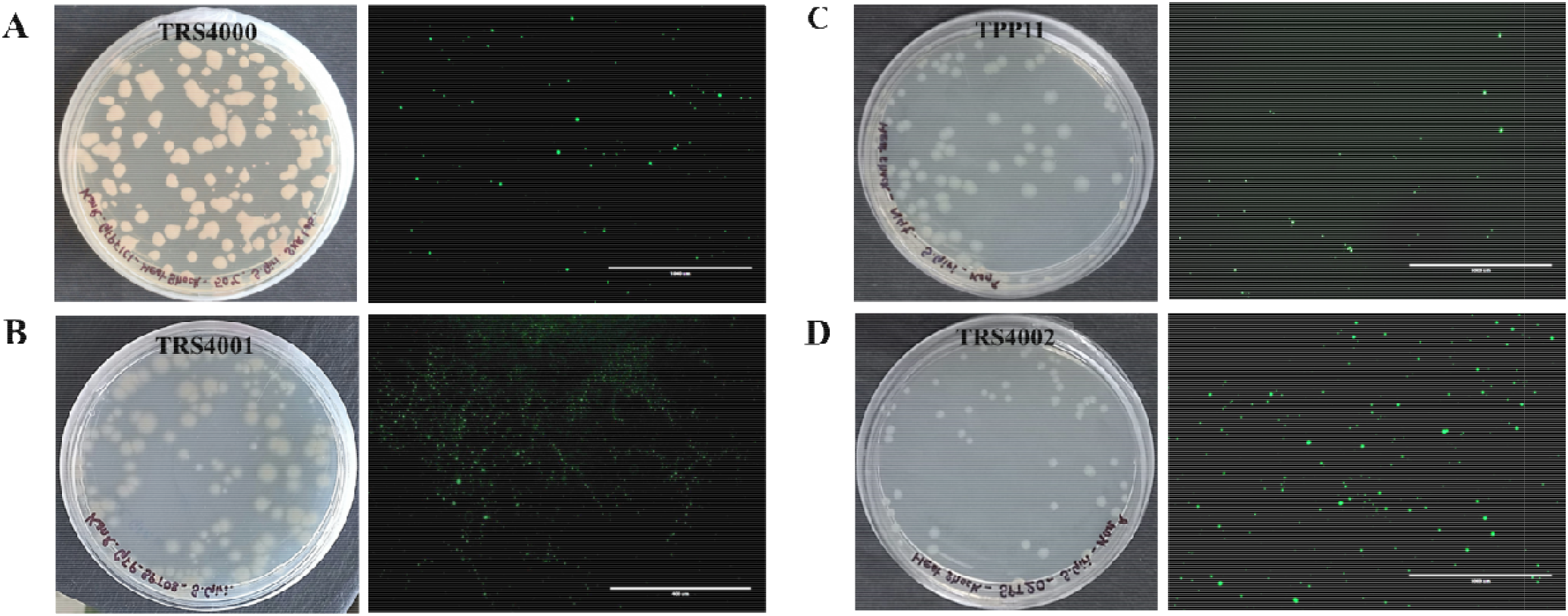
Heat shock-mediated transformation of GFP-plasmid (pDSK-GFPuv) in Gram-negative bacteria. Visualization of GFP-tagged bacterial isolates. The GFP-transformants were selected on BG agar plates using kanamycin (50 μg/mL) as a selection marker. The positive transformants *R. pseudosolanacearum* F1C1 (TRS4000) **(A)**, *P. aeruginosa* SPT08 (TRS4001) **(B)**, *P. putida* N4T (TPP11) **(C)**, and *E. roggenkampii* SPT20 (TRS4002) **(D)** cells containing the GFP plasmid on BG agar plate at 48 hours. All the strains were further grown in BG medium and observed under the fluorescence microscope using the GFP filter at 100X magnification with a scale bar of 400 μm.

### 3.2 Isolation and confirmation of GFP-plasmid from transformants

The plasmid (pDSK-GFPuv) was isolated from the transformants, which confirms the insertion of the plasmid into the targeted bacterial strains. The isolated plasmids were then treated with the enzyme *EcoRI* to confirm the size of the plasmids, which was found to be ∼ 8.9 kb, similar to the original plasmid that was kept as a positive control. Restriction analysis of plasmids confirmed the size and suggested that the plasmid doesn’t undergo any size changes after transformation. An uncut plasmid (pDSK-GFPuv) was kept as a positive control (Fig. 2A). The green fluorescence-tagged *R. pseudosolanacearum* F1C1 (TRS4000) was confirmed by phylotype-specific multiplex PCR, and wild-type F1C1 was taken as a positive control (Fig. 2B), confirming the transformation of the GFP-plasmid in the desired F1C1 strain. The plasmid (pDSK-GFPuv) was isolated from strain TRS4000, digested with *EcoRI*, which confirms the size of the plasmid that has been transformed (Fig. 2C and D). The transformants were characterized for twitching motility, where TRS4000 and TPA4001 show twitching motility activity, whereas TPP11 and TER4002 are twitching motility deficient, likewise the wild-type. The strain TRA4000 was found to be cellulase positive, and the other three bacterial strains were negative for the cellulase test, showing similar behavior to the wild type. *R. pseudosolanacearum* was inhibited by GFP-tagged strains TPA4001, TPP11, and TER4002, suggesting that the transformation of the GFP plasmid on the targeted bacterial strains couldn’t change the antibacterial activity of the bacterial strains (Fig. 2E). All the results confirmed the successful transformation of the GFP plasmid (pDSKGFPuv) into the targeted bacterial strains.

**Fig. 2.**
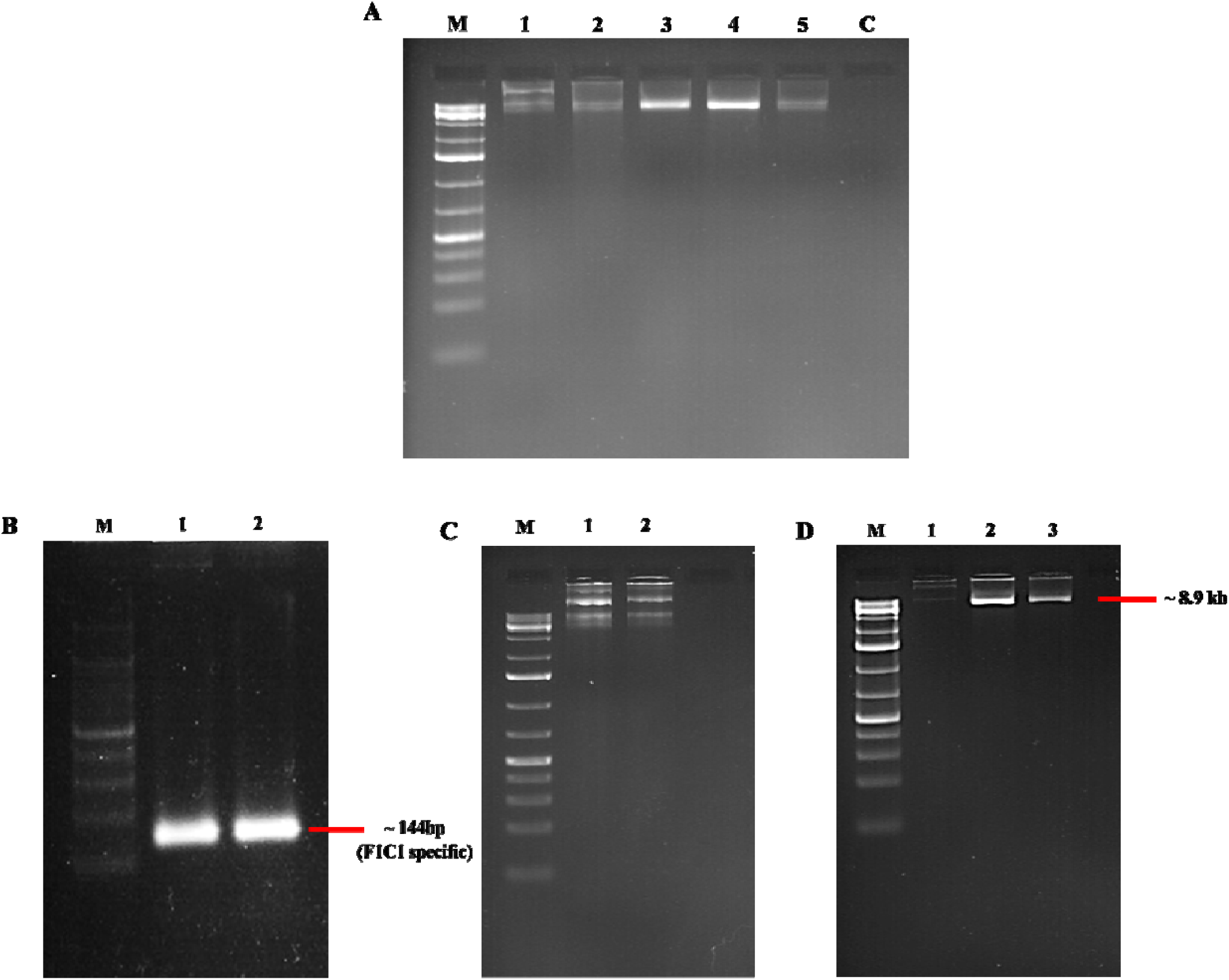
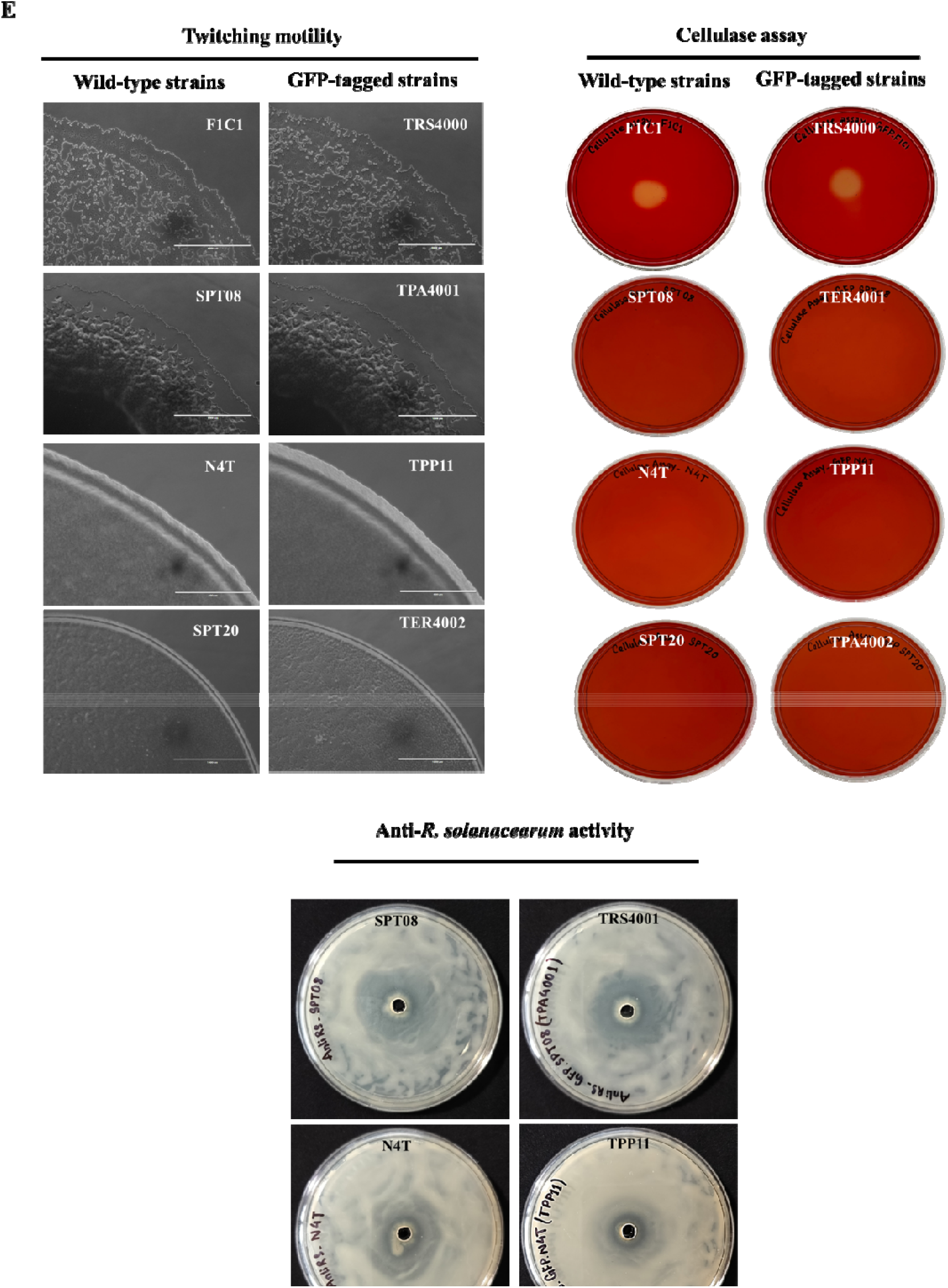
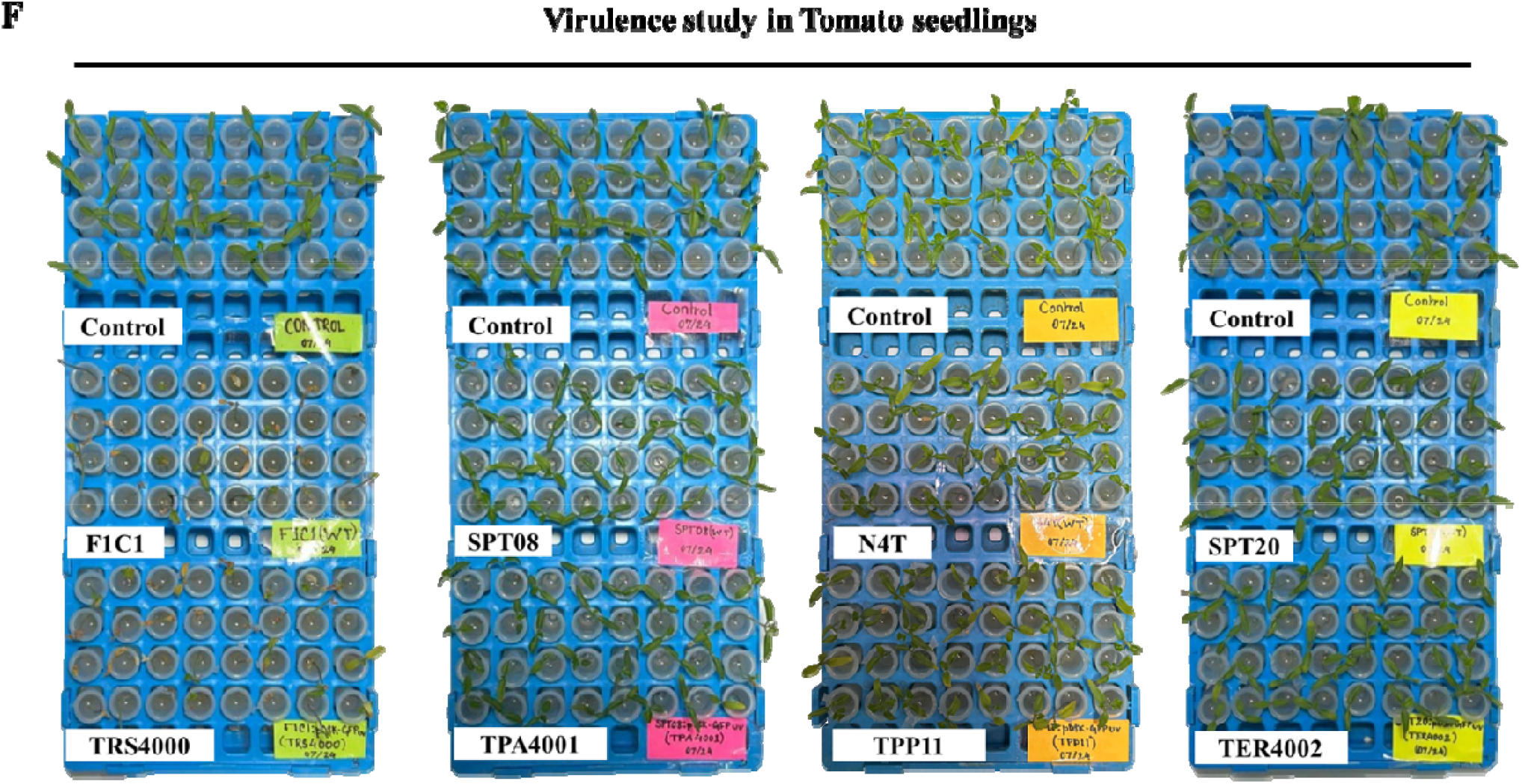
Confirmation of GFP-tagged transformants. Confirmation and characterization of GFP-tagged bacterial strains for successful transformation of green fluorescent plasmid (pDSK-GFPuv). Restriction analysis of pDSK-GFPuv plasmids isolated from the transformants (lane 2; uncut pDSK-GFPuv, lane 3; pDSK-GFPuv digested with *EcoRI*, lane 3, 4, 5; restriction digestion of pDSK-GFPuv isolated from TPA4001, TER4002, and TPP11) revealed the transformation (**A**), analyzed by gel electrophoresis on a 1% agarose gel. The GeneRuler 1 kb DNA marker (Thermo Scientific) is shown on the left (M). Phylotype-specific multiplex PCR of *R. pseudosolanacearum* F1C1 (lane 1) and GFP-tagged F1C1 (lane 2) confirmed successful transformation in the target strain (**B**). The 100 bp DNA marker was shown on the left (M). GFP-plasmid (pDSK-GFPuv) was isolated from strain TRS4000 (lane 2), and the original plasmid (lane 1), confirming plasmid insertion into the bacteria (**C**). Gel image showing the linearized pDSK-GFPuv-plasmid isolated from the strain TRS4000. The plasmid was digested with *EcoRI*, which yielded a single band at ∼ 8.9 kb of pDSK-GFPuv (lane 2; original plasmid, lane 3; TRS4000), and uncut pDSK-GFPuv (lane 1) was kept as a control (**D**). The GFP-tagged strains TRS4000, TPA4001, TPP11, and TER4002 were characterized by twitching motility, cellulase activity, and virulence assay, and compared with the wild-type strains (**E**). All GFP-tagged strains showed similar activity in all tests as the wild-type strains. Strains TRS4000 and TPA40001 exhibited twitching motility, while strains TPP11 and TER4002 lacked motility. All strains except TRS4000 were negative for the cellulase test, behaving similarly to the wild-type. The virulence study of strain TRS4000 showed it retained virulence similar to wild-type F1C1, while the other three strains had no negative impact on tomato seedlings (**F**). Virulence was assessed on one-week-old tomato seedlings via root inoculation and photographed at 10 days post-inoculation. The antibacterial activity of GFP-tagged strains of TPA4001, TPP11, and TER4002 was tested against *R. pseudosolanacearum* F1C1 to confirm their identity. F1C1 was inhibited by strains TPA4001, TPP11, and TER4002, indicating that plasmid transformation did not affect the antibacterial activity of the strains.

### 3.3 Study of plant-microbe association using GFP-tagged bacteria

The interaction of microbes within their hosts plays an important role in the survival of microbes within the plant. To study the bacterial interactions in the seedlings when observed under the microscope, the presence of a green fluorescence signal was observed after 24 hours in the root region, which represents the occurrence of bacteria inside the root. The fluorescence signal was observed in the root, stem, and leaf regions of the tomato seedlings after 24 hours of post-inoculation (Fig. 3), demonstrating their colonization patterns in the tomato seedlings.

**Fig. 3.**
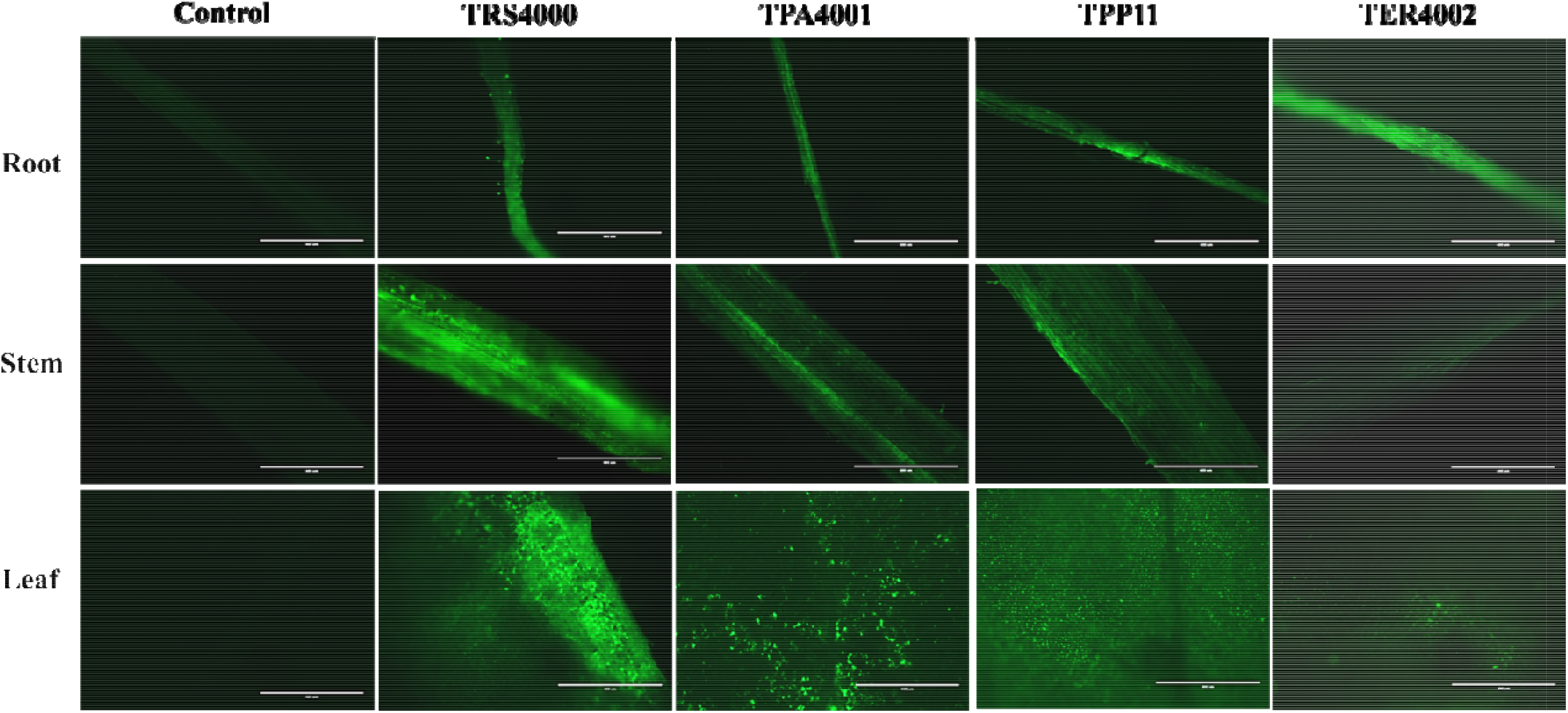
Localization of bacteria with GFP expression in tomato seedlings. Localization of GFP-tagged bacterial strains in one-week-old tomato seedlings. The bacterial strains TRS4000, TPA4001, TPP11, and TER4002 containing a GFP plasmid were inoculated into the one-week-old tomato seedlings by root inoculation. After 48 h of inoculation, a few of the seedlings were surface sterilized and observed under the fluorescence microscope (EVOS, Life Technologies). Tomato seedlings, such as the root, stem, and leaves, were observed with a fluorescence signal for each strain that was tagged with GFP, whereas in control seedlings inoculated with sterile water, no detectable signal was observed. All the photographs were obtained with 100X magnification with a scale bar of 400 μm. These bacterial strains, TPA4001, TPP11, and TER4002, were potential endophytes.

## 4 Discussion

Transformation is an important method to introduce external DNA into bacterial cells. Natural transformation, CaCl_2_-mediated heat shock transformation, and electroporation are the three techniques used by the researchers for transformation. Heat shock transformation is the most convenient one, though it is limited to smaller-sized DNA. In the present study, we have demonstrated the successful heat shock transformation in four different Gram-negative bacterial species, such as *Ralstonia pseudosolanacearum, Pseudomonas aeruginosa, Pseudomonas putida*, and *Enterobacter roggenkampii*. We believe that the result of this study will encourage other researchers to perform heat shock transformation in other bacteria.

*R. pseudosolanacearum* is an important plant-pathogenic bacterium. Both natural transformation and electroporation have been used to introduce external DNA into this bacterium with different success [27, 28]. The natural transformation procedure used for GMI1000 has not been found to be so consistent for studying the F1C1 strain, as the latter is found to grow in high concentrations of glycerol (unpublished results). The electroporation method used for transformation in this bacterium has limitations while studying GMI1000 strains [29]. The heat shock-mediated transformation has not been reported earlier for *Ralstonia solanacearum*. In this study, using the heat shock transformation technique, a green fluorescence gene-containing plasmid (∼8.9 kb) was transferred to the phytopathogen *R. pseudosolanacearum* F1C1, as well as three different plant-associated bacteria: *Pseudomonas aeruginosa* SPT08, *Pseudomonas putida* N4T, and *Enterobacter roggenkampii* SPT20. Although these four bacterial strains are Gram-negative, the transformation of the external plasmid was achieved at the same temperature with different times of heat treatment. The plasmid isolated from transformed F1C1 was again used to transform in *E. coli* to confirm its stability F1C1 (Fig. S1). Using this protocol, the plasmid has been successfully transformed into two other strains of *R. pseudosolanacearum* (JBF and BRS), which were isolated from banana (*Musa chinensis*) and brinjal (*Solanum melongena*), respectively (unpublished results from the laboratory). The heat shock-mediated transformation was also used to create a gene knockout mutation in the *R. pseudosolanacearum* F1C1 strain and complementation study using different plasmid (unpublished result from the laboratory).

To the best of our knowledge, this is the first demonstration of heat shock-mediated transformation in *Ralstonia pseudosolanacearum* using different plasmids. In addition, this protocol has been used to transfer the plasmid in *Pseudomonas* sp. and *Enterobacter* sp., suggesting that the heat shock transformation in Gram-negative bacteria establishes an efficient and straightforward protocol for transforming the GFP plasmid. Further research is required to evaluate the efficacy of this transformation method in bacteria using other vectors, plasmids, or constructs for bacterial transformation for its wider application.

## Author contributions

SJG wrote the draft, analyzed the data, reviewed, and edited. PLS, LD, and SB performed formal data analysis, reviewed, and edited the article. SB, MJ, and NA performed formal data analysis, reviewed the manuscript; SKR supervised, reviewed, edited, and conceptualized the work.

## Acknowledgements

We thank Prof. Kirankumar S. Mysore, Oklahoma State University, UAS, and former Professor, Plant Biology Division, The Samuel Roberts Noble Foundation, USA, for his kind gift of pDSK-GFPuv plasmid vector. Shubhra Jyoti Giri and Shuhada Begum are thankful for the JRF fellowship from the DBT, GoI, New Delhi grant (BT/PR41637/NER/95/1753/2021) awarded to Suvendra Kumar Ray. Pankaj Losan Sharma is thankful for the JRF/SRF fellowship from the DBT, GoI, New Delhi, grant (BT/403/NE/U-Excel/2013) awarded to Suvendra Kumar Ray. Lukapriya Dutta is thankful to DST, GoI, New Delhi, for the INSPIRE-SRF fellowship. Shuvam Bhuyan is thankful for the JRF/SRF fellowship from the UGC-NFSC, GoI, New Delhi, and DBT, GoI, New Delhi, for the Project Associate fellowship from the grant (BT/PR40253/BTIS/137/52/2022) awarded to Suvendra Kumar Ray. Monika Jain is thankful to the CSIR/JRF and UGC for the Non-NET fellowship.

## Declaration of completing interest

The authors declare that they do not have any conflict of interest.

## Ethical statement

No human participants and/or animals were used in the study.

## Notes

### Competing Interest Statement

The authors have declared no competing interest.

### Summary of Updates

The title has been modified; it was too long, and every bacterial strain was mentioned in the title, so we changed it

