## Supplementary Data for "Heat shock-mediated transformation is possible in several Gram-negative bacteria"

**Heat shock-mediated transformation in several Gram-negative bacteria such as *Ralstonia pseudosolanacearum*, *Pseudomonas aeruginosa*, *Pseudomonas putida,* and** ***Enterobacter roggenkampii***

Shubhra Jyoti Giri^1^, Pankaj Losan Sharma^1, 2^, Lukapriya Dutta^1^, Shuhada Begum^1^, Shuvam Bhuyan^1^, Monika Jain^1^, Niraj Agarwala^1, 3^, Suvendra Kumar Ray^1*^

^1^Molecular Plant-Microbe Interaction Laboratory, Department of Molecular Biology & Biotechnology, Tezpur University, Tezpur–784028, Assam, India.

^2^Department of Microbiology, Royal Global University, Guwahati-781035, Assam, India (Present Address)

^3^Department of Botany, Gauhati University, Guwahati-781014, Assam, India (Present Address)

**Figure legends**

**Figure 1.** **Transformation of pDSK-GFPuv isolated from *Ralstonia pseudosolanacearum* F1C1.** The GFP plasmid was isolated from the *R. pseudosolanacearum* F1C1, which was transformed into *Escherichia coli* DH5α cells **(B)** to ensure the plasmid transformation stability. The *E. coli* DH5α cells without the plasmid vector as a control (**A**).

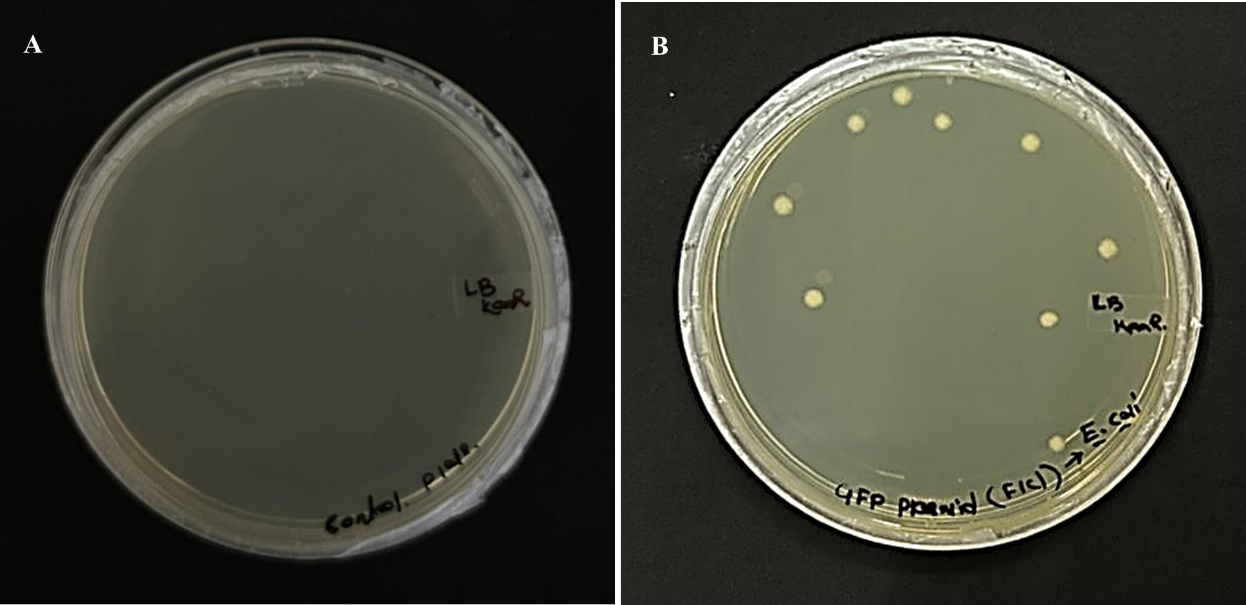
**Fig. 1**
